# Adaptive plasticity generates microclines in threespine stickleback male nuptial color

**DOI:** 10.1101/236943

**Authors:** Chad D. Brock, Molly E. Cummings, Daniel I. Bolnick

## Abstract

Adaptive phenotypic divergence is typically studied across relatively broad spatial scales (continents, archipelagos, river basins) because at these scales we expect environmental differences to be strong, and the homogenizing effect of gene flow to be weak. However, phenotypic plasticity and phenotype-dependent habitat choice are additional mechanisms that could also drive adaptation across spatially variable environments. We present evidence for apparently adaptive phenotypic variation across surprisingly small spatial scales (<2 vertical meters) in the threespine stickleback. We find that male breeding coloration varies as a function of the lakes’ optical-depth gradient, and these small-scale clines (‘microclines’) appear to be an adaptive response to ambient light gradients, as male color changes predictably in the opposite direction (‘countergradient’) to ambient light spectral shifts. Using visual models and field enclosure experiments, we show that these microclines result from phenotypic plasticity that maintains male conspicuousness. Our results show that adaptive phenotypic clines can exist across small spatial scales, because phenotypic plasticity rapidly generates repeatable trait-environment correlations despite the overwhelming opportunity for gene flow. Furthermore, these results provide strong evidence that phenotypic plasticity in nuptial coloration is an important mechanism for adjusting the conspicuousness of a visual signal to conspecifics.

## Introduction

Phenotype-environment correlations are often invoked as evidence for adaptation in response to spatially varying natural selection (Endler 1977; Langin et al. 2015; Richardson et al. 2014), especially when they are observed repeatedly in multiple independent populations (Hendry & taylor 2004; Colosimo et al. 2005; Wood et al. 2005; Langin et al. 2015). Theoretical work has shown that local adaptation can occur when selection is strong enough to overcome the homogenizing effect of gene flow, and empirical work has confirmed that migration load often constrains local adaptation (Felsenstein 1976; Garcia-Ramos & Kirkpatrick 1997).

Consequently, adaptive clines are typically studied at broad spatial scales at which gene flow is weak and environmental variation is pronounced (Hendry et al. 2002; Lenormand 2002; Richardson et al. 2014). However, alternative mechanisms, such as phenotypic plasticity (West-Eberhard 2003; Dewitt & Scheiner 2004; Engström-Öst & Candolin 2006; Lewandowski & Boughman 2008; Wund et al. 2008; Clarke & Schluter 2011; Svanback & Schluter 2012; Hendry 2016) and phenotype-dependent habitat choice (e.g. Edelaar et al. 2008; Bolnick et al. 2009) may also generate adaptive phenotypic divergence, including across much finer spatial scales than commonly assumed (Lenormand 2002; Richardson et al. 2014).

Male threespine stickleback (*Gasterosteus aculeatus*) exhibit diverse nuptial colors that females evaluate during mate choice (Boughman 2001). Male color varies among stickleback populations in a predictable manner (Reimchen 1989; Boughman 2001; Scott 2001). This variation is believed to be, at least partially, an adaptive response to the sensory environment, whereby male signals evolve to contrast against the optical background (Lythgoe 1979; Reimchen 1989; Endler 1992,1993; Boughman 2001; Scott 2001). For instance, in clear lakes the sidewelling ambient light is full spectrum and rich in blue wavelengths, which is thought to allow red-throated males to contrast strongly with their background. In tannic lakes where the sidewelling light is red-shifted, males tend to lose red throat coloration and instead develop a blue-black (“melanic”) phenotype (Reimchen 1989; Scott 2001). This color-environment correlation is thought to be an evolved response to divergent selection in allopatry or parapatry (Reimchen 1989; Boughman 2001; Scott 2001; Lewandowski & Boughman 2008; Malek et al. 2012; Yong et al. 2015). The repeatability of color variation suggests an adaptive value, whereas gene flow is weak or absent among populations.

Unexpectedly, we found these between-population trends recapitulated within lakes, where migration would typically be expected to erode any phenotype-environment correlations. In multiple lake populations of threespine stickleback we find microcline depth gradients in male nuptial color that are a predictable function of depth gradients in the optical environment. Consequently, the repeated, and ‘countergradient’ nature of the relationship between color and light environment suggests that male color signals are adapted to local nesting microenvironments, and our visual modeling analyses support this. Finally, we used field experiments and simulation models of stabilizing selection to provide strong support that phenotypic plasticity is the primary mechanism responsible for these microclines in male nuptial color.

## Methods

### Collections to test for male color microclines in naturally nesting stickleback

We investigated the relationship between stickleback male nuptial color and signaling environment within each of 15 lakes on Vancouver Island, British Columbia (*Supplementary Information 1*, Table S1). Males were collected directly from their nests by a snorkeler (CDB) using a dipnet, to ensure they were actively breeding.

For each nesting male collected, we recorded its nest depth, the distance to the nearest protective cover, and categorical descriptions of the substrate and nearby vegetation. Prior to capture, each male was observed for five minutes to note whether it displayed 1) aggression towards other stickleback, 2) courting behavior toward females, 3) nest-fanning behavior (a form of parental care). A darkened cooler with fresh lake water was used to transport males to shore for immediate collection of reflectance data (within 1 to 5 minutes after capture). This cooler was rinsed and refilled with lake water between males. Males were euthanized using MS-222 prior to the collection of reflectance data. <u>As male color can change rapidly after death</u>, the time of capture, euthanization <u>and reflectance measurements</u> were also recorded <u>for use as potential covariates in downstream analyses</u>. Furthermore, male and female color may vary throughout a breeding cycle (Hiermes et al. 2015, 2016). Consequently, we used our behavioral measurements as a proxy for male state (i.e. courting indicates a male is still receptive to gravid females) (see also Bolnick et al. 2015).

### Color measurement

For each male, spectral reflectance measurements were taken using a EPP200C UV-VIS spectrometer, SL-4 Xenon lamp, and a R400-7 reflectance probe for four body regions: 1) throat 2) pelvic region 3) preoperculum 4) lower iris. All measurements were taken while holding the probe perpendicular to the surface of the fish at a fixed 3mm distance. Three replicate measurements were collected for each body region, moving the probe slightly between replicates. Spectralon white standard measurements were taken between each fish to account for lamp drift.

We calculated the ratio of the respective areas under the reflectance curve for the wavelength intervals of 500-600 nm (green-orange) and 300-400 nm (UV-blue) for each body region. We focused on these two intervals because previous research has shown that both long wavelengths (e.g. orange and red) and short wavelengths (e.g. ultraviolet) strongly influence both mate preference and male-male aggressive encounters (Boughman 2001; Milinski & Bakker 1990; Rick & Bakker 2008a,b,c; Flamarique et al. 2013; Bolnick et al. 2016). Qualitatively similar results were found using the wavelength interval of 550-650 nm (orange-red) instead of the interval 500-600 nm. We next performed a principal components analysis on these ratios and extracted the first principal component as our color metric (PC1 variance explained=82.53%). The raw color ratios for all body regions loaded positively onto this first principal component axis, indicating that larger values of this first PC are associated with more reflectance in the green and orange range of the spectrum, relative to the UV and blue range, for all four body regions. Conversely, negative PC1 scores indicate relatively stronger UV and blue reflectance. We used the first PC axis as our focal measure of male color for all lakes except Gosling 2012 (see below), although qualitatively similar results were obtained using the ratio of green- and orange to UV- and blue reflectance for the preoperculum and abdomen separately. For the 60 experimental males collected in 2012 from Gosling Lake (see below), reflectance data were only collected from the preoperculum and abdomen regions, and we extracted the second principal component as our color metric (PC2 variance explained=19.74%). The raw color ratios for the preoperculum loaded positively on to this axis, while the abdomen color ratio loaded negatively. Equivalent results for these data were obtained using the raw color ratio of the preoperculum alone, though not the abdomen.

Preliminary analyses found no effect of behavior, nesting habitat (except for depth), or the time between collection, euthanization and reflectance measurement on male color, so we do not consider these further (see also Bolnick et al. 2015). Additionally, reflectance measurements for experimental males from Gosling Lake in 2012 were taken while alive (see below), and show similar results for both color and contrast to data collected in 2010 (when males were sacrificed prior to reflectance measurements). These results suggest that our subsequent analyses were not confounded by the effects of mortality-induced color change or variation due to reproductive state.

We fit a series of mixed models with our nuptial color metric as the response and nest depth as the only fixed predictor variable. Lake was also included as a random effect and the following nested models were compared using corrected Akaike Information Criterion (AICc) and likelihood-ratio tests (LRTs): 1) Random Intercept Only model 2) Nest Depth + Random Intercept model 3) Nest Depth + Random Slope + Random Intercept (Uncorrelated) 4) Nest Depth + Random Slope + Random Intercept (Correlated). Model (1) included no fixed effect of depth, and instead only allows for a different mean nuptial color for each lake. Model (2) includes nest depth as a fixed predictor, while also allowing for differences in mean nuptial color between lakes. Model (3) includes all terms from Model (2) as well as a random slope term that allows the relationship between color and depth to vary between lakes. Finally, Model (4) includes all terms from Model (3) with the addition of a correlation between the random intercept and random slope terms, which accounts for the possibility that redder or bluer lakes have predictably different depth gradients. As the assumption of a *χ*^2^-distributed test statistic is not met when testing borderline parameters such as random effects (e.g. testing whether σ_μ_=0), parametric bootstrapping (replicates=10000) was employed to calculate p-values for model comparisons. Additionally, within each lake we regressed the relevant PC axis versus nesting depth to estimate the male color depth gradient for comparison with the ambient light gradient (see below).

### Ambient light gradients

In each lake, we measured both sidewelling and downwelling irradiance along a depth gradient (0-2m) at 25-50 cm intervals. Irradiance data was collected with a EPP200C UV-VIS spectrometer and a UV-NIR cosine receptor. All irradiance data was collected in areas where males were found nesting that year. Five replicate measurements were taken at each depth and these measurements were dispersed throughout the nesting environment. We used the median value at each wavelength of these five replicates as our estimate of the ambient light at a given depth. As both the time of day and the time of year can impact irradiance measurements, all irradiance data was collected between 9-10:30 AM and within a window of 20 days (June 13 - July 2; this timeframe is in the middle of the breeding season of threespine stickleback in our study area). To control for ambient light conditions, we collected measurements above the water surface for each replicate to allow for the normalization of irradiance (Normalized Irradiance_Depth_=Irradiance_Depth_/Irradiance_Surface_). Additionally, we documented the ambient conditions at the time of each irradiance measurement (e.g. partly cloudy, sunny, etc.). Preliminary analyses indicated there was no significant effect of ambient condition on our irradiance measurements, and our study results were qualitatively similar regardless of whether we employed normalized or raw irradiance data. As such, we focus here on the results using only the latter.

Sidewelling irradiance was collected with the probe oriented horizontally away from the shore and represents the visual background against which a male stickleback is often viewed by females. Downwelling irradiance was measured with the probe directed vertically toward the water surface and represents the primary source of light for target reflection. We focus on sidewelling rather than downwelling irradiance because sidewelling light is the best predictor, by far, of among-population divergence in male color (Reimchen 1989; Boughman 2001; Scott 2001; Brock et al. *In preparation*). As with the male color data, we calculated the ratio of the respective areas under the spectral curve for the wavelength intervals of 500-600 nm (green-orange) and 300-400 nm (UV-blue) at each depth. Within each lake, we regressed this ratio versus depth to estimate the ambient light depth gradient. To compare ambient light and color-depth gradients we first used a sign test to assess whether gradient slopes were consistently associated across lakes in the predicted direction (i.e. color-depth gradients oriented in the opposite direction of the ambient light-depth gradients). Additionally, we employed weighted least squares regression to assess whether the magnitude, in addition to the orientation, of gradient slopes were correlated. For the latter analyses, we used the reciprocal of standard errors for each slope estimate as our weights. We applied these analyses to three datasets: 1) The full dataset (n=15 lakes) 2) all lakes with significant color-depth gradients (=color microclines), without a correction for multiple comparisons (n=9 lakes) 3) and finally the 6 lakes whose color-depth gradients survive multiple-test correction (FDR-corrected p<0.05).

### Stabilizing selection simulations

In principle, clinal variation in male color could arise within a single lake through a combination of natural selection and male site philopatry. To evaluate whether this selection-driven scenario is feasible, we used a simulation to estimate the strength of viability selection required to yield our observed microcline. We randomly generated a population of hypothetical male stickleback whose color (measured as the ratio of orange:UV) was drawn from a normal distribution whose mean and variance matched our 2012 Gosling Lake sample. We then randomly assigned each male to a nest depth from a uniform distribution whose range was dictated by the depth range in our 2012 Gosling Lake sample. No significant color-depth correlation resulted.We next simulated natural selection, as follows. We assumed that there is an optimal male color for a given nest depth, with stabilizing selection around that optimum such that an individual with color value *x* has fitness (Lande 1976):

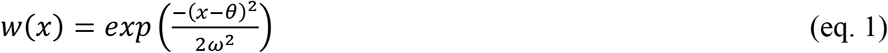

which is maximized at a fitness *w* = 1 when *x* = θ. The strength of stabilizing selection is inversely related to the width of the fitness landscape around θ, ω. We next assumed that θ varies linearly with male nest depth as follows:

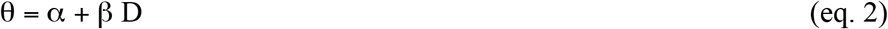

where *D* is nest depth and the parameters α and β are estimated from the slope and intercept of the empirical cline from Gosling Lake in 2012. Note that the microcline observed in 2012 was weaker than the 2010 microcline, so the following conclusions will be conservative in that even stronger selection would be necessary to generate the stronger 2010 trend. We then iterated through a range of values of ω (0.1 to 10 in increments of 0.1). For each value of ω, we randomly generated a population of 1000 males, randomly assigned them depths and color values as discussed above, then calculated their fitness given their depth and color, as defined in eqs. (1) and (2). A uniform random number was used to determine whether each male survived, with probability equal to *w*. The surviving males were then used to estimate a post-selection color-depth correlation. The fraction of non-survivors (mortality rate) was recorded as well, for all values of ω.

### Experimental test of color-dependent nest habitat choice

In June 2012 we constructed a 3m * 9m open-bottomed seine-net enclosure in Gosling Lake. The enclosure was set perpendicular to shore and spanned a depth gradient from 0.25m to 2.25m. The ambient light gradient was measured within the enclosure prior to the experiment. Using a Student’s t-test to compare the gradient slopes, we found that our experimental gradient did not differ significantly from the natural light-depth gradient (t=1.22, p=0.115).

All resident stickleback were removed from the enclosure, and it was restocked with twenty males and twenty females from the wild population in Gosling Lake. Prior to being introduced into the enclosure, reflectance data was collected from live males suspended in an aquarium constructed by CDB with UV-transmissive material. To minimize handling time, we collected data for only two body regions: the preoperculum and the pelvic region. To individually identify males, each was anesthetized using MS-222 and a coded wire tag (Northwest Marine Technologies, Shaw Island, WA, USA) was inserted under the skin anterior to the first dorsal spine prior to introduction into the enclosure. The enclosure was monitored daily by a snorkeler (CDB) to identify any males who had constructed nests and were actively courting females. These males were then captured and nest and reflectance data was collected as described above. We replaced any recaptured male with another wild-caught male (measured for color and tagged) to ensure that a fixed number of males (n=20) were always present within the enclosure. Similarly, any dead females or males were immediately removed from the enclosure and replaced by wild-caught fish. The experiment was terminated after 28 days.

Our first analysis was a linear regression to confirm that males used in this experiment exhibited a pre-experiment association between their initial color and their original nest depth. Next, we tested whether a male’s initial color predicted their subsequent choice of nest depth in the cage, using a linear regression in which depth is now the response variable and initial color is the predictor. Finally, we tested whether the original color-depth gradient was effectively recapitulated in the enclosure, by regressing a male’s final color on their chosen nest depth in the cage.

### Experimental test of phenotypic plasticity in male color

Concurrent with the habitat choice experiment, we tested whether males forced to nest at particular nest depths adjusted their color accordingly. We constructed 40 open-bottomed cylindrical enclosures (~1 m in diameter) out of 0.5 cm square hardware cloth, and installed these in Gosling Lake. We placed 20 enclosures in 0.45-0.55 m deep water, and the other 20 in 1.8 to 1.9 m deep water, with a minimum of 1 m of open space between each cage.

We placed one male and one gravid female into each cage at the start of the experiment. Gravid females were collected by trapping and dipnetting. Males were hand-collected from nests and their initial color was measured as described in the habitat choice experiment. Males and females were randomly distributed among the 40 cages. We placed a fixed amount of natural nesting material into each cage, which was replenished weekly. The experiment was run for 28 days to ensure the males had enough time to acclimate to these enclosures and, if possible, modify their color. Males were removed from cages in the same order in which they were added, and the total amount of time each male spent in an experimental enclosure was measured for use as a covariate.

We used linear regression to test whether initial male color was correlated with their original (natural) nest depth. A Student’s t-test was used to compare post-experiment male nuptial color between treatments. A linear mixed model (with male as a random effect) was used to assess whether the change in male color differed significantly between treatments and p-values for fixed effects were calculated using parametric bootstrapping (number of replicates=10000). As in the habitat choice experiment, we collected sidewelling irradiance for each enclosure prior to our experiment. We then compared the slope of the ambient light gradient of our enclosures to that of the lake as a whole using a Student’s t-test (t=1.12, p=0.137; *SI 7* & *8*, Fig. S2, S3). Finally, we tested whether the change in male color (Δcolor) depended on the magnitude and direction of change in nest depth (Δnest depth) using linear regression.

### Stickleback visual model to calculate chromatic contrasts

To assess whether the relationship between nuptial color and nest depth impacted male conspicuousness we calculated chromatic contrast, *ΔS*, using a stickleback-specific visual model. We developed a photoreceptor noise-limited color discrimination model for stickleback fish (Vorobyev & Osorio 1998) using MSP-estimated peak cone sensitivities (Rowe et al. 2004) and employed the parameters outlined in Govardovskii et al. (2000) to calculate spectral sensitivity functions for the four stickleback cone receptors (*SI 9*, Fig. S5). Detailed methods covering the construction and implementation of the visual model are given in *Supplementary Information 9*.

Chromatic contrast scores are calculated using the original reflectance data for a given body region. As the preoperculum is the only body region that is significantly related to nest depth across both years (2010 & 2012; See *Results*)) in Gosling Lake males, we chose to focus on this region when calculating chromatic contrast. To assess whether color shifts in our plasticity experiment lead to significant changes in *ΔS* across treatments we fit a general linear mixed model with the *ΔS* as the response and treatment and time (pre vs. post experiment) as fixed predictors as well as their interaction. Male stickleback ID was included as a random effect. As above, parametric bootstrapping (replicates=10000) was employed to calculate p-values for model terms.

Finally, to assess whether our color gradients were an artifact of the human sampling, we also developed a photoreceptor noise-limited color discrimination model for the human visual system based on data of human peak cone sensitivities from the literature (Bowmaker & Dartnell 1980). *ΔS* was calculated for a trichromatic visual system and linear regression and t-tests were employed to assess whether our sample was biased towards those individuals who were more conspicuous in their local nesting environment (see *SI 9* and Fig. S4 for methodological details and results).

## Results & Discussion

In a survey of 15 lake populations of threespine stickleback (*Supplementary Information 1* [*SI 1*], Table S1) we found that male color covaries with ambient light at fine spatial scales within populations. For example, in Gosling Lake male stickleback nest at depths ranging from 0.4 to 2.02 meters. Across this narrow depth gradient, the ambient light varies from UV-biased in shallows to orange-biased at depth. Consequently, deeper-nesting males reflect proportionally less green and orange and more UV and blue relative to shallow-nesting males, and this result was repeated across multiple years (Fig. 1b; 2010: n=13, r=-0.692, p=0.0088; 2012: n=60, r=-0.369, p=0.0037).

**Figure 1.**
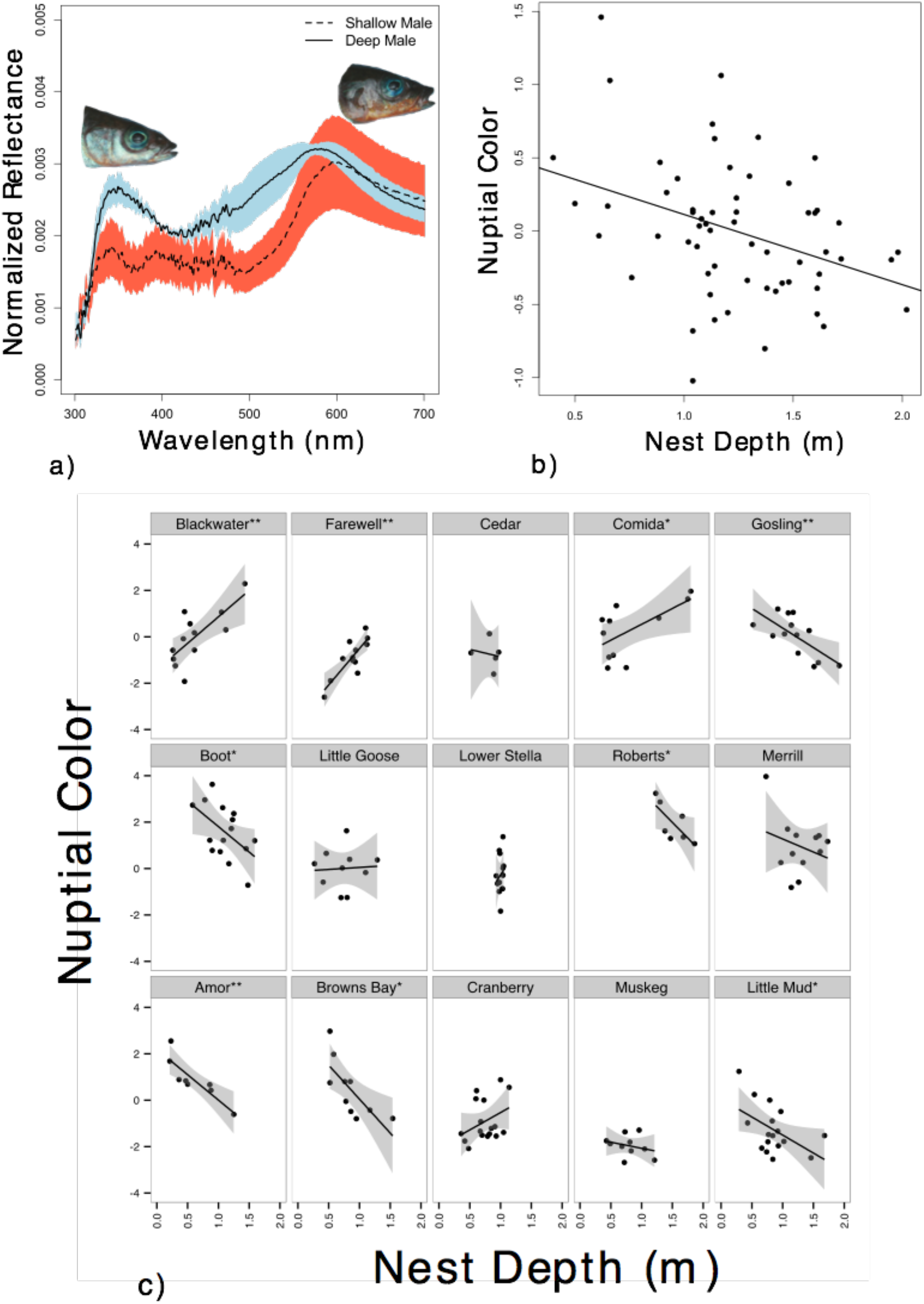
Color-depth microclines in threespine stickleback. **a)** Mean reflectance spectra of the preoperculum from males across two extremes of nest depth (0.4m and 2.02m for the shallow- and deep-nesting male, respectively) from a single location (Gosling Lake). Colored regions represent standard errors based on 3 replicate measurements for each male. **b)** Stickleback from Gosling Lake show a significant association between male nuptial color and nesting depth over spatial scales of <2 meters (n=60; r=0.369, p=0.004). **c)** Nine of 15 lakes show a significant relationship between color and nest depth before multiple-test correction (p<0.05; marked by single asterisks), while 6 remain significant after multiple-test correction (FDR-corrected p<0.05; marked by double asterisks). Shaded regions represent 95% confidence intervals for the slope of nuptial color ~ nest depth. Individual lake plots are ordered by their respective slope_irradiance_ from the most negative in the upper left to most positive in the lower right (*SI 2*).

This cline in Gosling Lake is not unique (and is not the result of human visual bias, *SI 9*, Fig. S4). We found significant male color ‘microclines’ in nine out of fifteen lakes examined (Fig. 1c; *SI 2*), always within a vertical range of less than 2 meters. The strength and orientation of these color-depth microclines varied among lakes (GLMM random slope effect; AIC_c weight_ =0.9957, *χ*^2^ =32.080, p < 0.0001; *SI 4*, Table S4; these results are robust to the exclusion of lakes with low depth coverage [<1m] for both color and irradiance, *SI 5*, Table S5). The repeatable presence of male nuptial color-depth correlations implies that they are non-random. We therefore hypothesized that male color gradients were an adaptive response to gradients in each lake’s optical environment.

Light wavelengths frequently change with water depth, but the direction of this change can depend on local dissolved tannin concentrations, suspended particles, and substrate composition (Lythgoe 1979; Endler 1992,1993). We confirmed that the optical background light (the Orange:UV proportion) changed with depth within each lake, but the slope of this gradient varied between lakes. Variation in optically-active substances means that some lakes will provide a more blue-shifted background at greater depth (negative slope_irradiance_) while others provide a red-shifted background at depth (positive slope_irradiance_, see *SI 2*, Table S2).

This among-lake variation in the direction of ambient light by depth gradients provides a unique opportunity to test sensory drive predictions regarding male nuptial coloration independent of habitat depth. Specifically, sensory drive theory (Endler 1992,1993) suggests that male color should negatively covary with the ambient light to maintain visibility to prospective mates. Consistent with sensory drive theory, we found that the color gradient slope negatively covaried with the ambient light gradients (sign test, p = 0.018; Weighted OLS regression, r=-0.46, p=0.07). This trend is also supported if we restrict our attention to the subset of lakes where color-depth gradients are significant before (9 lakes) and after (6 lakes) multiple test correction (9 lake subset: sign test, p = 0.019; Weighted OLS regression, r=-0.714, p=0.032; 6 lake subset, sign test, p = 0.016; Weighted OLS regression, r=-0.903, p=0.005). For example, in Gosling Lake the ambient light becomes more red-shifted at greater depths and males with deeper nests reflect more UV & blue (Fig. 1a,b, Fig. 2). Conversely, Blackwater Lake shows the opposite gradient in ambient light (i.e. background becomes more blue-shifted at greater depths), and males with deeper nests reflect more orange & red (Fig. 1c; *SI 2*). This predictable counter-gradient change in male color (in opposition to local sidewelling ambient light) provides strong evidence for an adaptive role of male signals in a changing optical environment (Fig. 2), consistent with sensory drive predictions.

**Figure 2.**
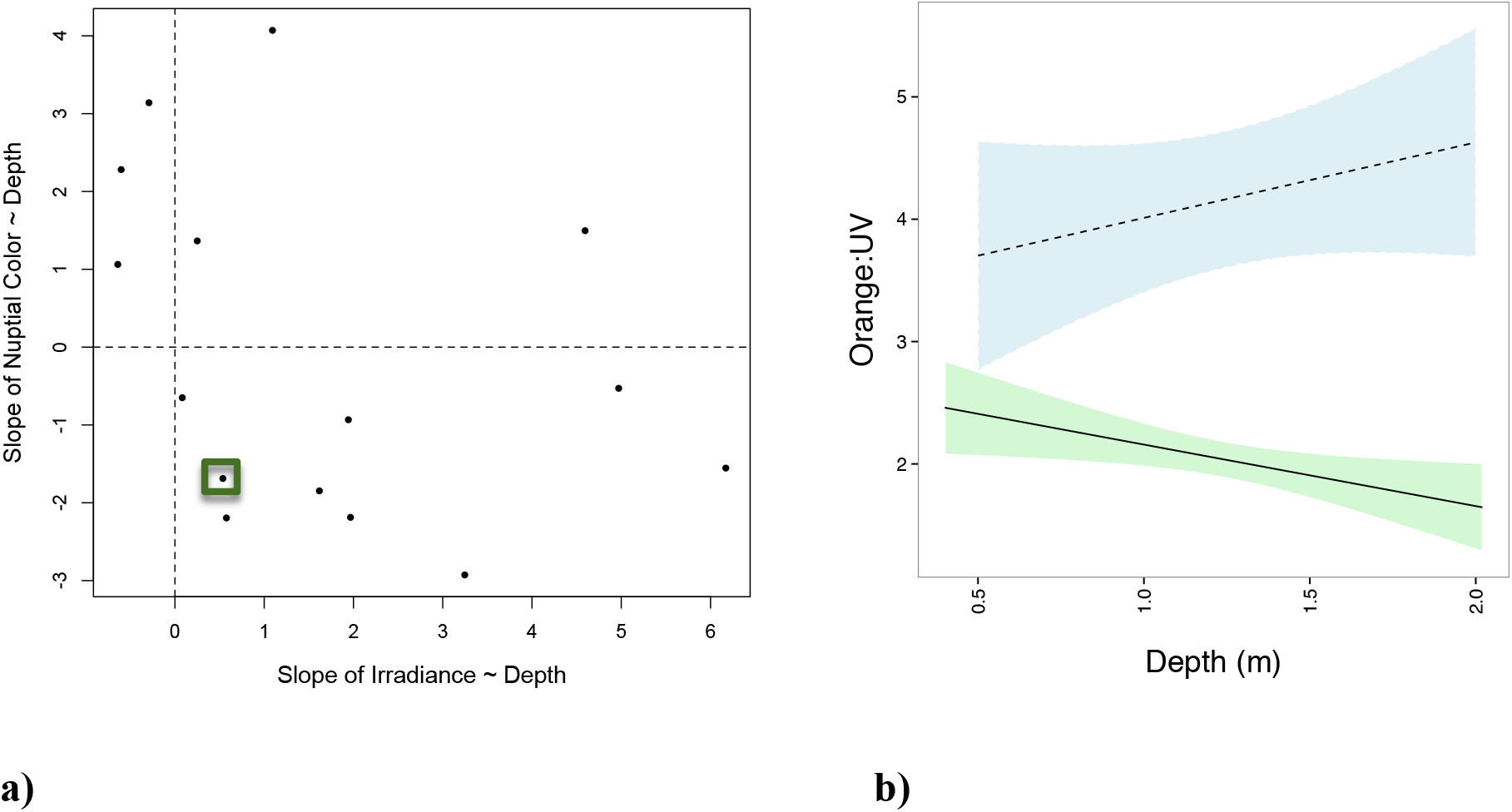
Color-depth microclines differ in a predictable manner across lakes. **a)** There is a predictable counter-gradient change in male color in opposition to local sidewelling ambient light in 12 of 15 lakes sampled as indicated by the significant relationship between the direction of the nuptial color-depth gradient (Slope_color_) and background light-depth gradient (Slope_irradiance_). Across 12 of our 15 lakes, the light background ~ depth gradient displays the opposite slope as the nuptial color-depth gradient as indicated by a sign test (15 lakes: p=0.018; 9 lakes: p=0.019; 6 lakes: p=0.016; See text for details on lake subsets). The box denotes the datapoint for Gosling Lake, which is plotted in detail in panel b). **b)** Plotted here is the line of best fit and 95% confidence intervals for the nuptial color ~ depth gradient (solid line) and the ambient background light ~ depth gradient (dashed line) in Gosling Lake for data collected in 2010.

As expected, our simulations demonstrate that stabilizing selection around a depth-specific optimal color is capable of generating a male color-depth microcline, assuming that males do not disperse (Fig. 4). Such a microcline would have to be re-established every breeding season if males choose their nest depth randomly, and this selection would have to act quickly to be observable in the first few weeks of a breeding season. More importantly, our simulations suggest that stabilizing selection would have to be exceptionally strong to generate the observed microcline<u>s</u>. Stabilizing selection in which 1/ω = 2.95 provides the best fit to the observed 2012 microcline (Fig. 4a). To illustrate how strong this selection must be, this value of 1/ω generated approximately 45% mortality (Fig. 4b) in order to yield the observed microcline. This mortality would have to take place within a few weeks following the initiation of breeding, given the timing of our empirical survey. Given this parameter estimate, quadratic regression of standardized fitness on standard normal color yields a quadratic selection gradient (γ) on the order of −1.6 which is far outside the range of typical empirical examples (Kingsolver et al. 2001; but see reference Stinchcombe 2008 for potential complications on interpreting Kingsolver et al.’s distribution of γ). While light environment can influence mortality rates in other fish (Maan et al. 2017), in our experimental transplant, which randomized males across nest depths much like in the simulation, we observed negligible mortality of males over a 4-week period (4 deaths out of 40 males=10% mortality) that did not differ significantly between treatments. We therefore conclude that natural selection is unable to account for the observed male microcline in Gosling Lake in 2012. The same inference applies to multiple other lakes, and to Gosling Lake in 2010 (when even stronger selection would have been required).

Two alternative mechanisms could give rise to microclines despite high dispersal rates. First, males might selectively choose nest depths that best suit a pre-existing phenotype (Edelaar et al. 2008; Bolnick et al. 2009). Alternatively, individuals may select a habitat at random (or based on an unrelated environmental variable), but subsequent phenotypic plasticity may alter their nuptial coloration to suit environmental conditions (West-Eberhard 2003; Dewitt & Scheiner 2004; McClennan 2007; Wund et al. 2008; Clarke & Schluter 2011; Svanback & Schluter 2012; Hendry 2016). We conducted two simultaneous experiments to distinguish these alternatives. First, we tested whether prior color influenced males’ choice of nest depth in a large (3*9m) rectangular cage spanning 0.25 to 2.25m depths in Gosling Lake, which has a positive irradiance-depth gradient (i.e. red-shifted at depth). Even though color was (again) negatively correlated with males’ original nest depth (n=20, r =-0.591, p=0.0061), this starting color did not influence males’ subsequent choice of nest depth within the cage (n=11, r=-0.217, p=0.635) and no color-depth correlation emerged given the opportunity for habitat choice (n=11, r=-0.358, p = 0.309). In the second (no-choice) study in Gosling Lake, males randomly assigned to nest in shallow water (~0.5 m) became more orange for all measured body parts (Fig. 3a), whereas males in deep cages (~1.85 m) became bluer (difference in final color: *t* = −3.2882, p = 0.0027; Time*Treatment effect in a GLMM: p = 0.0382). Males subjected to the greatest change in nest depth exhibited the largest change in color, though this result was only marginally significant (r=-0.327, p = 0.05; Fig. 3c). Furthermore, the experimentally induced plastic changes in male color resulted in a color-depth slope that accounted for most (~70%; *SI 7*, Fig. S2) of the naturally observed microcline slope. This demonstrates that plasticity is a viable mechanism for maintaining adaptive coloration across various (or varying) optical environments in stickleback fish. A stickleback visual model revealed that the experimentally induced color changes in Gosling Lake increased the chromatic contrast (*ΔS*) of deep-water males, but, surprisingly, decreased the contrast for shallow-water males (GLMM Time*Treatment p=0.0002; Fig. 3b). This functional inference recapitulates the tendency for chromatic contrast to increase with nest depth in wild males (Brock et al., *In press)*. We speculate that this depth gradient in contrast may reflect selection from aerial predators for reduced visibility near the surface, as has been demonstrated in some cichlid fish (Maan et al. 2008), or selection for traits in addition, or even contrary to, those that increase conspicuousness (Gong & Gibson 1996; Brock et al., *In press*).

**Figure 3.**
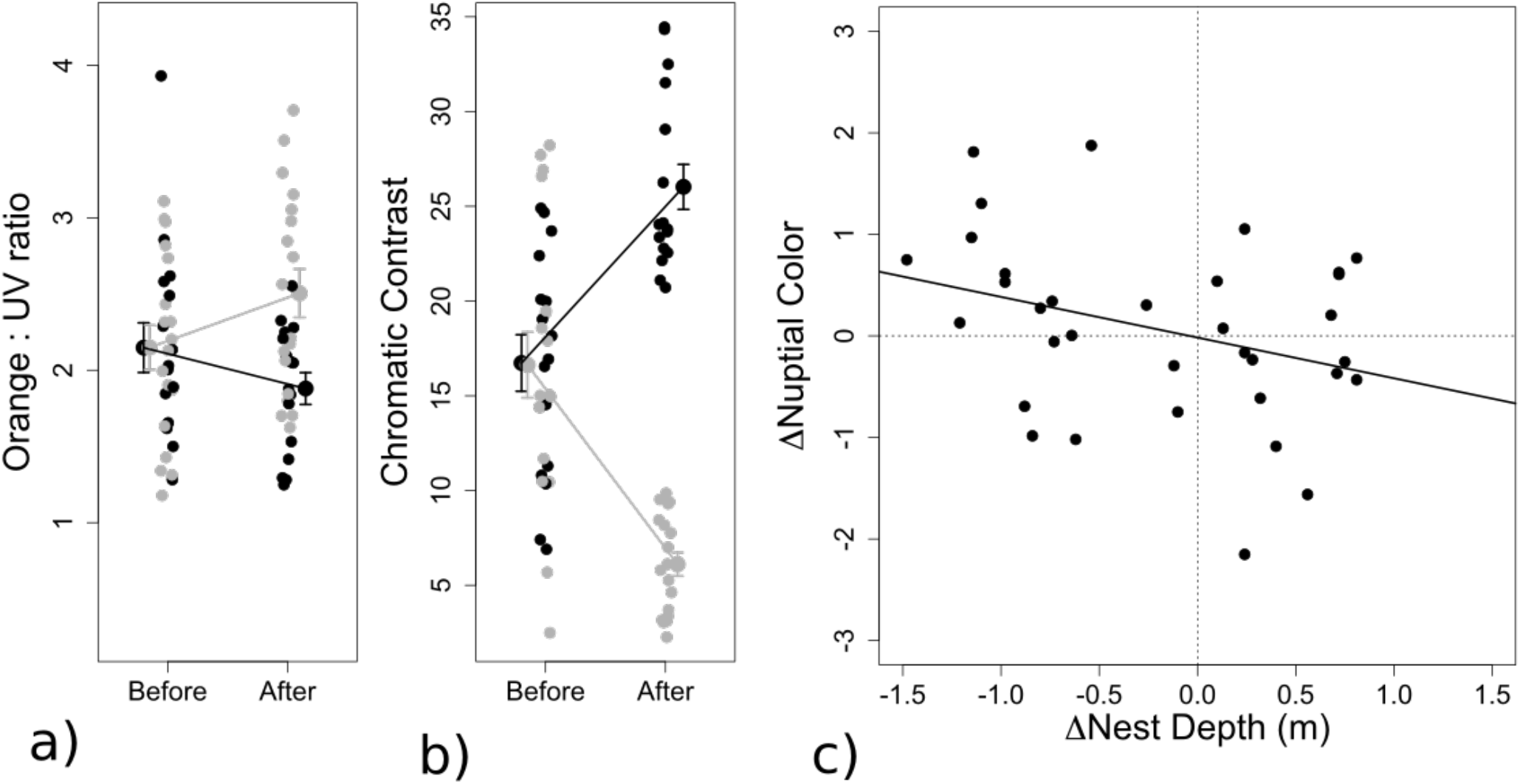
Phenotypic plasticity leads to color-depth microclines. **a)** Male nuptial color changes in a predictable fashion across depth treatments. As in wild caught fish, the ratio of orange:UV was on average higher in fish from shallow enclosures (light gray) than fish from deep enclosures (black) (*t*=-3.2882, p=0.0027). This was due primarily to a differential shift in color between treatments (GLMM Time*Treatment p=0.0382). **b)** Alterations in male nuptial color over the 4-week experimental period lead to extreme differences in chromatic contrast (*ΔS*) between depth treatments (GLMM Time*Treatment p<0.001). **c)** More drastic changes in nest depth are associated with larger changes in nuptial color, though this relationship is not significant (p=0.05).

**Figure 4.**
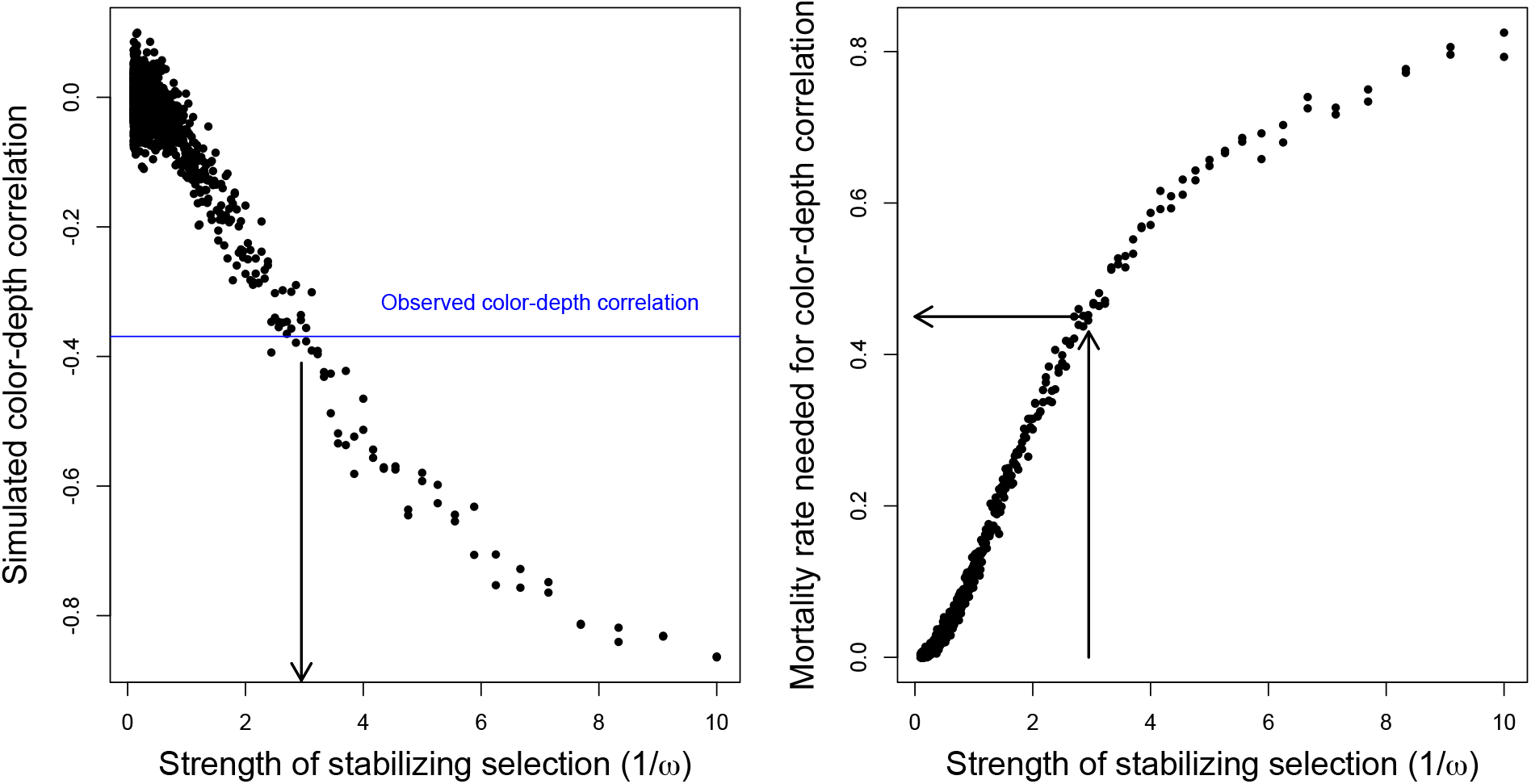
Evaluating the capacity of natural selection to yield observed microclines. In the simulation, we assumed that males randomly pick a nest depth, and then viability selection acts to eliminate males whose color is poorly matched to their assigned depth, resulting in a simulated color-depth correlation among survivors. We then identified the strength of viability selection necessary to generate our observed correlation, and evaluated whether it is empirically reasonable. **a)** The strength of stabilizing selection (arrow) that results in a color-depth correlation that matches the empirical correlation from wild caught Gosling males in 2012 (blue line). **b)** The mortality rate associated with the strength of stabilizing selection necessary to recapitulate the wild color-depth microclines (arrows).

The nature and/or the intensity of predation has been shown to vary across habitat depth in a number of freshwater fish species (Power 1984; Maan et al. 2008; but see Rypel et al. 2007 & Bossu and Near 2015). In threespine stickleback, Reimchen (1980) provided evidence for spatial variation in predation mode, though not intensity, across limnetic vs. littoral environments, though how this relates to water depth is unknown (But see Doucette et al. 2004; Brown & Moyle 1997). Preliminary data from GoPro cameras in Gosling Lake suggest that trout and bellastomatid density vary across water depths, though these patterns are not significant (Brock et al. *In press*). Consequently, the role that predation plays in this system remains an open question. More extensive data on predator distribution and abundance across lakes and nesting habitats, and the employment of predator-specific visual models (e.g. Crothers & Cummings 2013) could help elucidate the role that predation plays in generating these male color microclines.

Other potential factors may also influence male color and contrast across nest depths, including microspatial variation in diet (Boughman 2007; Snowberg & Bolnick 2012), immune function (Kurtz et al. 2007; Bolnick et al. 2015b), stickleback density (Candolin & Salesto 2009), female preference (Bolnick et al. 2015a), and the strength of sexual selection (Bolnick et al. 2015a; Bolnick et al. 2016). Furthermore, as male color varies with health and condition (e.g. Milinski & Bakker 1990; Frischknecht 1993; Boughman 2007), direct and/or indirect benefits to female mate choice may drive preference toward hues that are more strongly correlated with these factors, even if these hues are less conspicuous (Brock et al. *In press*).

## Conclusion

Our data demonstrate that stickleback male nuptial color shows predictable microclinal variation across nesting depth. Although depth gradients in fish color have been shown before (Seehausen et al. 2008; Marques et al. 2017), the stickleback microclines span a smaller spatial scale than previously reported (<2 meter vertical range in depth despite relatively high stickleback mobility). The repeatable and predictable nature of these male nuptial color microclines implies that they are an adaptation to micro-spatial variation in optical environments. Specifically, both wild and experimentally caged stickleback exhibit nuptial colors that differ from the dominant wavelength of the background light. An exceptional feature of our data is the fact that ambient light gradients differ among lakes, providing an unprecedented test of sensory drive theory. We find that male clines are consistently counter-gradient to ambient light. Deeper-nesting males tended to exhibit more chromatic contrast against their background light, which should make them particularly conspicuous to females. Shallow-nesting males were relatively cryptic, potentially decreasing their probability of being detected by surface or littoral predators (Maan et al. 2008). This fitting of male nuptial color to optical environment confirms a key expectation of sensory drive theory (Reimchen 1989; Endler 1992, 1993; Boughman 2001; Scott 2001; Fuller 2002; Cummings 2007; Seehausen et al. 2008; Ryan & Cummings 2013). Finally, although selection and habitat choice may play a role in generating microclines, our experiments in Gosling Lake clearly show that phenotypic plasticity is nearly sufficient to generate the microcline on a short time-scale.

These conclusions have several broad implications. First, microspatial clines may help explain a long-standing puzzle in evolutionary genetics: the maintenance of genetic and phenotypic variation within natural populations. Many populations inhabit landscapes in which biotic and abiotic variables change subtly over fine spatial scales, possibly leading to micro-spatial variation in selection, which in turn can facilitate the maintenance of polymorphisms, including diverse color phenotypes (Chunco et al. 2007; Gray & Mckinnon 2007; Gray et al. 2008; Marques et al. 2017). Second, our results highlight the role of phenotypic plasticity in generating repeatable and thus presumably adaptive clines. While previous work has shown male color is plastic in threespine stickleback (e.g. Engström-Öst & Candolin 2006; Lewandowski & Boughman 2007; Clarke & Schluter 2011), to our knowledge our study provides the first evidence for plasticity driving repeatable and presumably adaptive associations between color and optical environment. The influence of plasticity on phenotypic evolution remains an intriguing open question in this system and others (Ancel 2000; Price et al. 2003; West-Eberhard 2003; Dewitt & Scheiner 2004; Engström-Öst & Candolin 2006; Ghalambor et al. 2007; Paenke et al. 2007; Wund et al. 2008; Ghalambor et al. 2015; Hendry 2016). However, given evidence from previous studies that nuptial color differences between stickleback populations have a genetic component (Lewandowski & Boughman 2008; Malek et al. 2012; Yong et al. 2015), our results suggest that plasticity could play a significant role in facilitating phenotypic and genetic divergence across greater spatial and temporal scales (Price 2008; Wund et al. 2008; Pfennig et al. 2010; Fitzpatrick 2012; Rebar & Rodriguez 2014a,b). Finally, prior studies suggest that depth-mediated light gradients can contribute to speciation via sensory drive in fish such as cichlids, which exhibit depth gradients (across 4-8m) in both male nuptial color and opsin pigments (Seehausen et al. 2008). Stickleback fish exhibit appreciable assortative mating within populations (e.g. Snowberg & Bolnick 2008; Bolnick et al. 2015a), as well as depth gradients in opsin expression (Veen et al. *In press*). This raises the intriguing but as-yet-untested possibility that microgeographic variation in male nuptial color might represent an initial step in the evolution of reproductive isolation (Boughman 2001; Marques et al. 2017). Such reproductive isolation would act, at least initially, on microclines arising mostly from phenotypic plasticity, an often-overlooked force in species formation (Price 2008; Wund et al. 2008; Pfennig et al. 2010; Fitzpatrick 2012; Rebar & Rodriguez 2014a,b).

## Acknowledgements

We thank Dale Jacques, Chris Smith, Jason Lu, Kim Ballare, Cathy Hernandez, and Jesse Weber for assistance with field collections, and Mike Ryan, Mark Kirkpatrick, Hans Hofmann and the Wagner Lab for their comments on the manuscript. The research was supported by NSF grant IOS-1145468 to DIB, a David and Lucille Packard Foundation Fellowship (DIB), and the Howard Hughes Medical Institute (DIB).

## Author contributions

CDB, DIB & MJC designed the study, CDB conducted the study and collected the data. CDB, DIB & MJC conducted the analyses and wrote the paper.

## Materials & correspondence

All correspondence and requests for materials should be directed to CDB at.

## References

Ancel, L. W. (2000) Undermining the Baldwin expediting effect: does phenotypic plasticity accelerate evolution? Theor. Popul. Biol. 58: 307–319.

Bolnick, D. I. and P. Nosil. (2007) Migration load and the strength of selection. Evolution, 61: 2229–2243.

Bolnick, D. I. et al. (2009) Phenotype-dependent native habitat preference facilitates divergence between parametric lake and stream stickleback. Evolution 63: 2004–2016.

Bolnick, D. I., K. C. Shim, M. Schmerer, & C. D. Brock. 2015a. Population-specific covariation between immune function and color of nesting male threespine stickleback. PLoS One. 10: e0126000. do:10.1371/journal.pone.0126000.

Bolnick, D. I., K. C. Shim, & C. D. Brock. 2015b. Female stickleback prefer shallow males: sexual selection on nest microhabitat. Evolution 69 1643–1653.

Bolnick, D. I., K. Hendrix, L.A. Jordan, T. Veen & C. D. Brock. 2016. Intruder colour and light environment jointly determine how nesting male stickleback respond to simulated territorial intrusions. Biol. Lett. 12: 20160467.

Boughman, J. W. (2001) Divergent sexual selection enhances reproductive isolation in sticklebacks. Nature, 411: 944–948.

Bossu, C. M. & T. J. Near. 2015. Ecological constraint and the evolution of sexual dichromatism in darters. Evolution. 69 1219–1231.

Boughman, J. W. 2001. Divergent sexual selection enhances reproductive isolation in sticklebacks. Nature, 411 944–948.

Boughman, J. W. 2007. Condition dependent expression of red color differs between stickleback species. J. Evol. Biol. 20: 1577–1590.

Bowmaker, J. K. & H. J. Dartnall. (1980) Visual pigments of rods and cones in a human retina. J. Physiol. 298: 501–511.

Candolin U. & T. Salesto 2009. Does competition allow male mate choosiness in threespine stickleback? Am. Nat. 173 273–277.

Chunco, A. J., J. S. McKinnon & M. R. Servedio. (2007) Microhabitat variation and sexual selection can maintain male colour polymorphisms. Evolution. 61: 2504–2515.

Clarke, J. M. and D. Schluter. (2011) Colour plasticity and background matching in a threespine stickleback species pair. Biol. J. Linn. Soc. 102: 902–914.

Colosimo, P. F. et al. (2005) Widespread parallel evolution in sticklebacks by repeated fixation of ectodyplasin alleles. Science, 307: 1928–1933.

Crothers L.C. & Cummings M.E. 2013. Warning signal brightness variation: sexual selection may work under the radar of natural selection in populations of a polytypic poison frog. Am. Nat. 181(5): E116–E124. doi: 10.1086/670010.

Cummings, M. E. (2007) Sensory trade-offs predicts signal divergence in surfperch. Evolution. 61: 530–545.

Dewitt, T. J. and S. M. Scheiner (Eds.). Phenotypic plasticity: functional and conceptual approaches. (Oxford Univ. Press 2004).

Doucette, L. I., S. Skulason & S. S. Snorrason. 2004. Risk of predation as a promoting factor of species divergence in threespine sticklebacks (*Gasterosteus aculeatus* L.). Biol. J. Linn. Soc. 82 189–203.

Edelaar, P., A. M. Siepelski & J. Clobert. (2008) Matching habitat choice causes directed gene flow: a neglected dimension in evolution and ecology. Evolution. 62: 2462–2472.

Endler, J. (1993) Some general comments on the evolution and design of animal communication systems. Philos. Trans. R. Soc. Lond. Ser. B. 340: 215–225.

Endler, J. A. (1992) Signals, signal conditions, and the direction of evolution. Am. Nat. 139: 125–153.

Endler, J. A. Geographic Variation, Speciation, and Clines (Princeton University Press 1977).

Engstrom-Ost, J. & U. Candolin (2006). Human induced water turbidity alters selection on sexual displays in stickleback. Behav. Ecol. 59: 689–693.

Felsenstein, J. The theoretical population genetics of variable selection and migration. (1976) Ann. Rev. Genet. 10: 253–280.

Fitzpatrick, B. M. (2012) Underappreciated consequences of phenotypic plasticity for ecological speciation. Int. J. Ecol. 2012:256017 doi:10.1155/2012/256017.

Flamarique, I. N., C. Bergstrom, C. L. Chang & T. E. Reimchen (2013) Role of the iridescent eye in stickleback female mate choice. J. Exp. Biol. 216: 2806–2812.

Frischknecht, M. 1993. The breeding coloration of male threespined sticklebacks (*Gasterosteus aculeatus*) as an indicator of energy investment vigour. Evol. Ecol. 7: 439.

Fuller, R. C. (2002) Lighting environment predicts relative abundance of male color morphs in bluefin killifish populations. Proc. R. Soc. Lond. B, 269: 1457–1465.

Garcia-Ramos, G. and M. Kirkpatrick. (1997) Genetic models of adaptation and gene flow in peripheral populations. Evolution 51: 21–28

Ghalambor, C. K. et al. (2015) Non-adaptive plasticity potentiates rapid adaptive evolution of gene expression in nature. Nature. 525: 372–375.

Ghalambor, C. K., McKay, J. K., Carroll, S. P. & Reznick, D. N. (2007) Adaptive versus non-adaptive phenotypic plasticity and the potential for contemporary adaptation in new environments. Funct. Ecol. 21: 394–407.

Gong, A. and R. M. Gibson. (1996) Reversal of a female preference after visual exposure to a predator in the guppy, Poecilia reticulata. Anim. Behav. 52: 1007–1015.

Govardovskii, V. I., N. Fyhrquist, T. Reuter, D. G. Kuzmin, K. Donner. (2000) In search of the visual pigment template. Vis. Neuro. 17: 509–528.

Gray S. M. & J. S. McKinnon. (2007) Linking colour polymorphism maintenance and speciation. TREE 22: 71–79

Gray, S. M., L. M. Dill, F. Y. Tantu, E. R. Loew, F. Herder & J. S. McKinnon. (2008) Environment-contongent sexual selection in a color polymorphic fish. Proc. Roy. Soc. B. 275: 1785–1791.

Hendry, A. Eco-evolutionary Dynamics. (Princeton University Press 2016).

Hendry, A. P et al. (2002) Adaptive divergence and the balance between selection and gene flow: lake and stream in the misty system. Evolution, 56: 1199–1216.

Hendry, A. P. and E. B. Taylor. (2004) How much of the variation in adaptation can be explained by gene flow? An evaluation using lake-stream stickleback pairs. Evolution, 58: 2319–2331.

Kingsolver J. G. et al. (2001) The Strength of Phenotypic Selection in Natural Populations. Evolution. 157 245–261.

Kirk, J. T. O. Light and photosynthesis in aquatic ecosystems. (Cambridge Univ. Press 1994).

Kurtz, J., M. Kalbe, A. Langefors, I. Mayer, M. Milinski, and D. Hasselquist. (2007) An experimental test of the immunocompetence handicap hypothesis in a teleost fish: 11-Ketotestosterone suppresses innate immunity in three-spined sticklebacks. Am. Nat. 170: 509–519.

Lande, R. (1976) Natural selection and random drift in phenotypic evolution. Evolution. 30 314–334.

Langin, K. M., T. S. Sillett, W. C. Funk, S. A. Morrison, M. A. Desrosiers, and C. K. Ghalambor. (2015) Islands within an island: Repeated adaptive divergence in a single population. Evolution 69: 653–665.

Lenormand, T. (2002) Gene flow and the limits to natural selection. Trends Ecol. Evol., 17: 183–189.

Lewandowski, E. & J. Boughman. (2008) Effects of genetic and light environment on colour expression in threespine stickleback. Biol. J. Linn Soc. 94: 663–673.

Lythgoe, J. N. The Ecology of Vision. (Oxford Univ. Press 1979).

Malek, T. B., J. W. Boughman, I. Dworkin, C. L. Peichel. (2012) Admixture mapping of male nuptial color and body shape in a recently formed hybrid population of threespine stickleback. Mol. Ecol. 21: 5265–5279.

Marques, D. A., K. Lucek, M. P. Haesler, A. F. Feller, J. I. Meier, C. E. Wagner, L. Excoffier & O. Seehausen. (2017) Genomic landscape of early ecological speciation initiated by selection on nuptial colour. Mol. Ecol. 26: 7–24.

McLennan, D.A. The Umwelt of the Threespine Stickleback. In: Ostlund-Nilsson S., Mayer, I., Huntingford, F.A. editors. Biology of the Three-spined Stickleback. CRC Press; pp. 179–211. (2007)

Milinski, M. & T. C. M. Bakker (1990) Female sticklebacks use male coloration in mate choice and hence avoid parasitized males. Nature. 344: 330–333.

Maan, M. et al. (2008) Color polymorphism and predation in a Lake Victoria cichlid fish. Copeia, 3: 621–629.

Paenke, I., Sendhoff, B. & Kawecki, T. J. (2007) Influence of plasticity and learning on evolution under directional selection. Am. Nat. 170: E47–E58.

Pfennig, D. W. et al. (2010) Phenotypic plasticity’s impact on diversification and speciation. TREE. 25: 459–467.

Power, M. E. 1984. Depth distributions of armored catfish: predator-induced resource avoidance? Ecology. 65 523–528.

Price, T. D., Qvarnström, A. & Irwin, D. E. (2003) The role of phenotypic plasticity in driving genetic evolution. Proc. R. Soc. Lond. B 270: 1433–1440.

Price, T. Speciation in Birds. (Roberts & Company Publishers 2008).

Rebar, D. & R. L. Rodriguez. (2014) Genetic variation in host plants influences the mate preferences of a plant-feeding insect. Am. Nat. 184: 489–499.

Rebar, D. & R. L. Rodriguez. (2014) Trees to treehoppers: genetic variation in host plants contributes to variation in the mating signals of a plant-feeding insect. Ecol. Lett. 17: 203–210.

Reimchen, T. E. (1989) Loss of nuptial color in threespine sticklebacks (Gasterosteus aculeatus). Evolution, 43: 450–460.

Richardson, J. L., M. C. Urban, D. I. Bolnick, and D. K. Shelly. (2014) Microgeographic adaptation and the spatial scale of evolution. Trends Ecol. & Evol. 29: 165–176.

Rick, I. P. & T. C. M. Bakker (2008a) Color signaling in conspicuous red sticklebacks: do ultraviolet signals surpass others? BMC Evol. Biol. 8: 189

Rick, I. P. & T. C. M. Bakker (2008b) Males do not see only red: UV wavelengths and male territorial aggression in the three-spined stickleback (*Gasterosteus aculeatus*). Naturwissenschaften. 95: 631–638.

Rick, I. P. & T. C. M. Bakker (2008c) UV wavelengths make female three-spined sticklebacks (*Gasterosteus aculeatus*) more attractive for males. Behav. Ecol. Sociobiol. 62: 439–445

Rick, I. P., D. Bloemker & T. C. M. Bakker (2011) Spectral composition and visual foraging in the three-spined stickleback (Gasterosteidae: *Gasterosteus aculeatus* L.): elucidating the role of ultraviolet wavelengths. Biol. J. Linn. Soc. 105(2): 359–368.

Rowe, M. P., C. L. Baube, E. R. Loew, and J. B. Phillips. (2004) Optimal mechanism for finding and selecting mates: how threespine stickleback (Gasterosteus aculeatus) should encode male throat colors. J. Comp. Physiol. A. 190: 241–256

Ryan M. J. and M. E. Cummings. (2013) Perceptual biases and mate choice. Ann. Rev. Ecol. Evol. Sys. 44: 437–459.

Rypel, A.L., C.A. Layman and D.A. Arrington. 2007. Water depth modifies relative predation risk for a motile fish taxa in Bahamian tidal creeks. Estuaries and Coasts, 30: 518–525.

Sabbah, S. et al. (2011) The underwater photic environment of Cape Maclear, Lake Malawi: comparison between rock- and sand-bottom habitats and implications for cichlid fish vision. J. Exp. Biol. 214: 487–500.

Scott, R. J. (2001) Sensory drive and nuptial color loss in the three-spine stickleback. J. Fish Biol. 59: 1520–1528.

Seehausen, O. et al. (2008) Speciation through sensory drive in cichlid fish. Nature. 455 620–626.

Snowberg, L. K. and D. I. Bolnick. (2008) Assortative mating by diet in a phenotypically unimodal but ecologically variable population of sticklebacks. Am. Nat. 172: 733–739.

Snowberg, L.S., and D.I. Bolnick. (2012) Partitioning the effects of spatial isolation, nest habitat, and individual diet in causing assortative mating within a population of threespine stickleback. Evolution. 66: 3582–3594. doi: 10.1111/j.1558-5646.2012.01701.x PMID: 23106720

Stinchcombe, J. R. et al. (2008) Estimating nonlinear selection gradients using quadratic regression coefficients: double or nothing? Evolution. 62 2435–2440.

Svanbäck, R. and D. Schluter. (2012) Niche specialization influences adaptive phenotypic plasticity in threespine stickleback. Am. Nat. 180: 50–59.

Vorobyev, M. and D. Osorio. (1998) Receptor noise as a determinant of colour thresholds. Proc. Roy. Soc. B. 265: 351–358.

West-Eberhard, M. J. Developmental Plasticity and Evolution (Oxford Univ. Press, 2003).

Wood, T. E., J. M. Burke, and L. H Rieseberg. (2005) Parallel genotypic adaptation: when evolution repeats itself. Genetica, 123: 157–170.

Wund M. et al. (2008) A test of the “Flexible Stem” model of evolution: Ancestral plasticity, genetic accommodation, and morphological divergence in the threespine stickleback radiation. Am. Nat. 172(4): 449–462.

Yong, L., C. L. Peichel & J. S. McKinnon. (2015) Genetic architecture of conspicuous red ornaments in female threespine stickleback. Bethesda. 6: 579–588.

